# A robust dissociation among the language, multiple demand, and default mode networks: evidence from inter-region correlations in effect size

**DOI:** 10.1101/140384

**Authors:** Zachary Mineroff, Idan Blank, Kyle Mahowald, Evelina Fedorenko

**Affiliations:** Massachusetts Institute of Technology; Harvard Medical School; Massachusetts General Hospital

**Keywords:** functional MRI, language, multiple demand network, default mode network, individual-subject analyses

## Abstract

Complex cognitive processes, including language, rely on multiple mental operations that are carried out by several large-scale functional networks in the frontal, temporal, and parietal association cortices of the human brain. The central division of cognitive labor is between two fronto-parietal bilateral networks: (a) the multiple demand (MD) network, which supports executive processes, such as working memory and cognitive control, and is engaged by diverse task domains, including language, especially when comprehension gets difficult; and (b) the default mode network (DMN), which supports introspective processes, such as mind wandering, and is active when we are not engaged in processing external stimuli. These two networks are strongly dissociated in both their functional profiles and their patterns of activity fluctuations during naturalistic cognition. Here, we focus on the functional relationship between these two networks and a third network: (c) the fronto-temporal left-lateralized “core” language network, which is selectively recruited by linguistic processing. Is the language network distinct and dissociated from both the MD network and the DMN, or is it synchronized and integrated with one or both of them? Recent work has provided evidence for a dissociation between the language network and the MD network. However, the relationship between the language network and the DMN is less clear, with some evidence for coordinated activity patterns and similar response profiles, perhaps due to the role of both in semantic processing. Here we use a novel fMRI approach to examine the relationship among the three networks: we measure the strength of activations in different language, MD, and DMN regions to functional contrasts typically used to identify each network, and then test which regions co-vary in their contrast effect sizes across 60 individuals. We find that effect sizes correlate strongly within each network (e.g., one language region and another language region, or one DMN region and another DMN region), but show little or no correlation for region pairs across networks (e.g., a language region and a DMN region). Thus, we replicate the language/MD network dissociation discovered previously with other methods, and also show that the language network is robustly dissociated from the DMN, overall suggesting that these three networks support distinct computations and contribute to high-level cognition in different ways. Inter-individual differences in effect sizes therefore do not simply reflect general differences in vascularization or attention, but exhibit sensitivity to the functional architecture of the brain. The strength of activation in each network can thus be probed separately in studies that attempt to link neural variability to behavioral or genetic variability.

## Introduction

High-level cognition is supported by the frontal, temporal, and parietal association cortices, which have vastly expanded in the human brain compared to the brains of our closest primate relatives (e.g., Buckner and Krienen, 2013). These cortices are not organized into individual, “isolated” regions, but rather consist of multiple large-scale “networks”: sets of regions that share structural and functional properties (e.g., Alexander-Bloch et al., 2013; Bernard et al., 2012; Chen et al., 2012; Crossley et al., 2013; de Pasquale et al., 2012; Fox et al., 2005; Golland et al., 2007; Hagmann et al., 2008; Kalcher et al., 2012; Konopka et al., 2012; Power et al., 2011; Raznahan et al., 2011; Seeley et al., 2009; Toro et al., 2008; van den Heuvel and Sporns, 2011; Vértes et al., 2012; Wu et al., 2011; Yeo et al., 2011). The central division of cognitive labor is between two such networks: (a) the fronto-parietal bilateral multiple demand (MD) network (e.g., Duncan, 2010, 2013; Duncan and Owen, 2000; see also Braver et al., 2003; Cabeza and Nyberg, 2000; Cole and Schneider, 2007; Dosenbach et al., 2007, among others), which supports diverse goal-directed behaviors (e.g,. Fox et al., 2005; Stiers et al., 2010) and is modulated by general cognitive effort (e.g., Duncan and Owen, 2000; Fedorenko et al., 2013; Hugdahl et al., 2015); and (b) the fronto-parietal bilateral default mode network (DMN; Andrews-Hanna et al., 2010; Buckner et al., 2008; Humphreys et al., 2015; Raichle et al., 2001), which supports more “restful”, internally-oriented, processes (e.g., Gusnard and Raichle, 2001; Raichle et al., 2001), such as mind-wandering, reminiscing about the past, and imagining the future (e.g., Buckner et al., 2008; Spreng et al., 2009). The distinct and complementary functions of these two networks have long been recognized (e.g., Fox et al., 2005; Fransson, 2005; Golland et al., 2007; Greicius et al., 2003; Uddin et al., 2009).

However, these networks are not the only contributors to complex cognition in the associative cortices, and the functional relationship between each of these two networks and other high-level networks is less clear. Here, we focus on the relationship between the MD and default-mode networks and a third network: the fronto-temporal left-lateralized “core” language network (e.g., Fedorenko et al., 2010), which selectively supports language processing (e.g., Fedorenko et al., 2011; Monti et al., 2012). Is this network distinct and dissociated from the MD network and the DMN, or is it functionally synchronized and integrated with one or both of them? One might hypothesize that the language network is integrated, to some extent, with the MD network given that language processing requires general attention, working memory, and cognitive control (e.g., see Fedorenko, 2014; Novick et al., 2010, for reviews). And one might also hypothesize that the language network is integrated with the DMN given that a lot of our introspective processing plausibly draws on verbal resources (e.g., Carruthers, 2002; Morin and Michaud, 2007; Pléh, 2002; Schrauf, 2002; Sokolov, 1972; Vygotsky, 1962, 2012; Zivin, 1979).

Until recently, researchers have actually not explicitly distinguished between the language and the MD networks, especially in the frontal lobes, where subsets of each network reside side by side within the region known as “Broca’s area” (Fedorenko et al., 2012). However, recent work has established that these networks are spatially and functionally distinct based on two converging lines of evidence. First, language and MD regions exhibit *distinct functional profiles*: whereas MD regions are recruited across many cognitive tasks, language regions respond selectively during language processing and are not engaged by a wide range of non-linguistic processes, including arithmetic, working memory, cognitive control, and music perception (e.g., Fedorenko et al., 2011; Monti et al., 2012; see Fedorenko and Varley, 2016, for a recent review). And second, language and MD regions show *distinct patterns of fluctuations in neural activity* during naturalistic cognition. For example, Blank et al. (2014, replicated in Paunov et al., submitted) compared fluctuations in the fMRI BOLD signal across language and MD regions either during “rest” or while participants listened to stories. In both conditions, the average pairwise correlations among language regions (see also Hampson et al., 2002; Turken and Dronkers, 2011; Yue et al., 2013) and among MD regions (see also Dosenbach et al., 2007; Hampshire et al., 2012; Seeley et al., 2007) were significantly higher than correlations between pairs of regions straddling the two networks, which were close to zero. This dissociation was further supported by data-driven clustering of BOLD signal time-courses, which grouped language and MD regions separately.

The relationship between the language network and the DMN remains less clear. For example, algorithms that cluster regions across the brain based on similarities in their activity fluctuations during rest often produce a cluster whose topography resembles a union of the language network and the DMN (e.g., Yeo et al., 2011). However, interpreting the resulting cluster in functional terms can be difficult (see e.g., Blank and Fedorenko, submitted; Blank et al., 2014 for a discussion) and must rely on logically flawed “reverse inference” from anatomical coordinates back to cognitive processes (Poldrack, 2006, 2011). Furthermore, at least some of the language regions appear to deactivate during some demanding cognitive tasks (e.g., Fedorenko et al., 2011; see also Figure 2, top panel), which is one functional signature of the DMN – although, unlike DMN regions, language regions increase their activity during difficult language processing tasks (e.g., Blank et al., 2016). Finally, both language and DMN regions have been linked to semantic / conceptual processing (e.g., Binder et al., 2009; Jackson et al., 2016; Wirth et al., 2011). However, damage to each network produces distinct behavioral deficits: deficits in language comprehension and production for the language network (e.g., Bates et al., 2003; Mesulam et al., 2015; Mirman et al., 2015; Ojemann et al., 2003), and deficits in e.g., autobiographical memory retrieval for the DMN (e.g., Damasio and Van Hoesen, 1983; Philippi et al., 2015).

To shed further light on the relationship among the language, multiple demand, and default mode networks, here we characterize and directly compare their functional properties using fMRI. We first examine the *basic response profiles* of language, MD, and DMN regions – defined functionally in each of 60 individual participants – and show that the profiles are clearly distinct. We then use a novel approach to probe the relationship among the three networks, testing whether *the strength of functional responses in these three networks co-varies* across participants. This approach is inspired by several recent findings. First, different language regions robustly co-vary across individuals in their respective effect sizes for a contrast between reading sentences and reading lists of nonwords (Mahowald and Fedorenko, 2016). Second, different MD regions co-vary across individuals in their respective effect sizes for a contrast between hard and easy spatial working memory task (Assem et al., submitted). Importantly, these effect sizes appear to be highly stable within participants across runs and scanning sessions, suggesting that they tap some time-invariant idiosyncratic properties of individual brains. Here, we extend this study of effect-size correlations from pairs of regions within a single network to pairs of regions across different networks. If these effect size measures reflect some highly general functional properties, like the degree of brain vascularization or fluid intelligence levels, then all three networks should co-vary in these measures across individuals. However, if these measures are sensitive to functional dissociations among distinct brain networks, we expect the language and MD networks to show little co-variation in these measures across participants, consistent with prior studies (e.g., Blank and Fedorenko, submitted; Blank et al., 2014; Fedorenko et al., 2011; Fedorenko et al., 2012; Paunov et al., submitted). Critically, if effect size measures indeed respect such functional distinctions, then the degree to which the language and DMN regions co-vary across individuals could indicate the extent of functional association between these two networks. To foreshadow our conclusions, these measures replicate the robust language-MD dissociation, and show that the language network and the DMN are also robustly dissociated.

## Methods

### Participants

Sixty participants (41 females) between the ages of 19 and 45 – students at MIT and members of the surrounding community – were paid for their participation. Participants were right-handed native speakers of English, naïve to the purposes of the study. All participants gave informed consent in accordance with the requirements of MIT’s Committee On the Use of Humans as Experimental Subjects (COUHES).

### Design, materials, and procedure

Each participant performed two tasks that were designed to localize the functional networks of interest: a reading task for the language network (adapted from Fedorenko et al., 2010) and a spatial working memory (WM) task for the MD network and DMN (from Fedorenko et al., 2011). Some participants also completed one or two additional tasks for unrelated studies. The scanning session lasted approximately 2 hours.

#### Language localizer task

Participants read sentences (e.g., *NOBODY COULD HAVE PREDICTED THE EARTHQUAKE IN THIS PART OF THE COUNTRY*) and lists of unconnected, pronounceable nonwords (e.g., *U BIZBY ACWORRILY MIDARAL MAPE LAS POME U TRINT WEPS WIBRON PUZ*) in a blocked design. Each stimulus consisted of twelve words/nonwords. For details of how the language materials were constructed, see Fedorenko et al. (2010). The materials are available at http://web.mit.edu/evelina9/www/funcloc/funcloc_localizers.html. The sentences > nonword-lists contrast has been previously shown to reliably activate high-level language processing regions and to be robust to the materials, task, and modality of presentation (Fedorenko et al., 2011; Fedorenko et al., 2010; Mahowald and Fedorenko, 2016; Scott et al., 2016).

Stimuli were presented in the center of the screen, one word/nonword at a time, at the rate of 450ms per word/nonword. Each stimulus was preceded by a 100ms blank screen and followed by a 400ms screen showing a picture of a finger pressing a button, and a blank screen for another 100ms, for a total trial duration of 6s. Participants were asked to press a button whenever they saw the picture of a finger pressing a button. This task was included to help participants stay alert and awake.

Condition order was counterbalanced across runs. Experimental blocks lasted 18s (with 3 trials per block), and fixation blocks lasted 14s. Each run (consisting of 5 fixation blocks and 16 experimental blocks) lasted 358s. Each participant completed 2 runs.

#### Spatial working memory task

Participants had to keep track of four (easy condition) or eight (hard condition) sequentially presented locations in a 3×4 grid (Fedorenko et al., 2011). In both conditions, participants performed a two-alternative forced-choice task at the end of each trial to indicate the set of locations they just saw. The hard > easy contrast has been previously shown to robustly activate MD regions (Blank et al., 2014; Fedorenko et al., 2013). The reverse contrast, easy > hard, robustly activates DMN regions, in line with prior work using similar tasks and contrasts (Fedorenko et al., 2013; Leech et al., 2011; McKiernan et al., 2003; Park et al., 2010).

Stimuli were presented in the center of the screen across four steps. Each of these steps lasted for 1000ms and presented one location on the grid in the easy condition, and two locations in the hard condition. Each stimulus was followed by a choice-selection step, which showed two grids side by side. One grid contained the locations shown on the previous four steps, while the other contained an incorrect set of locations. Participants were asked to press one of two buttons to choose the grid that showed the correct locations.

Condition order was counterbalanced across runs and participants. Experimental blocks lasted 32s (with 4 trials per block), and fixation blocks lasted 16s. Each run (consisting of 4 fixation blocks and 12 experimental blocks) lasted 448s. Each participant completed 2 runs.

### fMRI data acquisition

Structural and functional data were collected on the whole-body, 3 Tesla, Siemens Trio scanner with a 32-channel head coil, at the Athinoula A. Martinos Imaging Center at the McGovern Institute for Brain Research at MIT. T1-weighted structural images were collected in 176 sagittal slices with 1mm isotropic voxels (TR=2530ms, TE=3.48ms). Functional, blood oxygenation level dependent (BOLD), data were acquired using an EPI sequence (with a 90° flip angle and using GRAPPA with an acceleration factor of 2), with the following acquisition parameters: thirty-one 4mm thick near-axial slices acquired in the interleaved order (with 10% distance factor), 2.1mm×2.1mm in-plane resolution, FoV in the phase encoding (A>>P) direction 200mm and matrix size 96mm×96mm, TR=2000ms and TE=30ms. The first 10s of each run were excluded to allow for steady state magnetization.

### fMRI data preprocessing

MRI data were analyzed using SPM5 and custom Matlab scripts (available in the form of an SPM toolbox from http://www.nitrc.org/projects/spm_ss). Each participant’s data were motion corrected and then normalized into a common brain space (the Montreal Neurological Institute (MNI) template) and resampled into 2mm isotropic voxels. The data were then smoothed with a 4mm FWHM Gaussian filter and high-pass filtered (at 200s). Effects were estimated using a General Linear Model (GLM) in which each experimental condition was modeled with a boxcar function (modeling entire blocks) convolved with the canonical hemodynamic response function (HRF).

### Defining individual functional regions of interest (fROIs)

All the analyses described below were performed on the responses in regions of interest that were defined functionally in each individual participant (e.g., Fedorenko et al., 2010; Nieto-Castañón and Fedorenko, 2012; Saxe et al., 2006). Three sets of functional regions of interest (fROIs) were defined, one set for each of the three networks. To do so, we used the Group-constrained Subject-Specific (GSS) approach developed in Fedorenko et al. (2010; Julian et al., 2012). In particular, fROIs were constrained to fall within a set of “masks” which marked the expected gross locations of activations for the relevant contrast (based on prior work). For the language network, the masks were generated based on a group-level representation of data from 220 participants (see Figure 1; these masks are similar to the masks originally reported in Fedorenko et al., 2010 based on 25 participants, except that the left anterior temporal and left mid-anterior temporal masks are grouped together, and the left mid-posterior temporal and left posterior temporal masks are grouped together). In addition, given that both the MD network and the DMN are bilateral, we defined the RH homologs of the LH language regions by transposing the LH masks onto the RH, as in Blank et al. (2014). fROIs were then defined in each RH mask, and were allowed the differ from their left homologues in their precise locations within these masks. For the MD and DMN networks, the masks were mostly anatomical regions from the Tzourio-Mazoyer et al. (2002) atlas, selected based on the prior literature (see Figure 1; the anatomical MD masks were the same as those used in Fedorenko et al., 2013 and Blank et al., 2014). The only exceptions are the bilateral Temporo-Parietal Junction masks for the DMN, which were created based on a random-effects analysis of activation maps for a functional contrast from a Theory of Mind localizer (false belief > false photograph) from 462 participants (Dufour et al., 2013).

**Figure 1.**
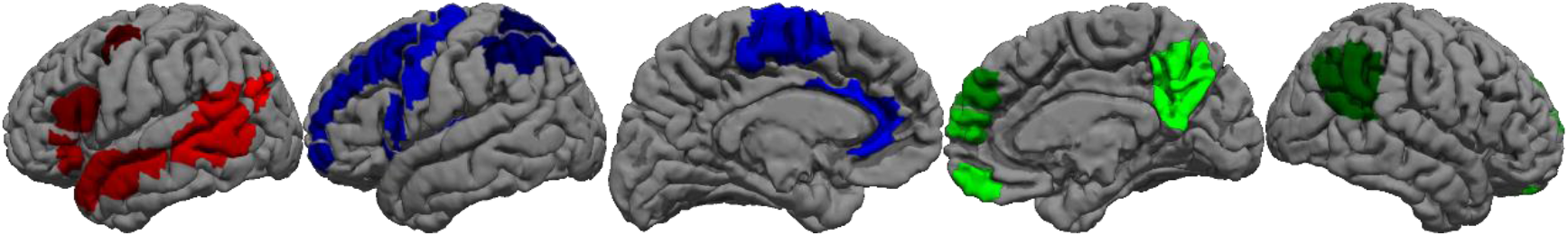
The “masks” for the language (red), multiple demand (blue), and default mode (green) networks used to constrain the selection of individual functional ROIs.

For each participant, each set of masks was intersected with their individual activation map for the relevant contrast (i.e., sentences > nonwords for the language network, hard > easy spatial WM for the MD network, and easy > hard spatial WM for the DMN). Within each mask, the voxels were then sorted based on their *t* values for the relevant contrast, and the top 10% of voxels were selected as that participant’s fROI. This top *n*% approach ensures that the fROIs can be defined in every participant – thus enabling us to generalize the results to the entire population (Nieto-Castañón and Fedorenko, 2012) – and that fROI sizes are the same across participants.

For the language network, twelve fROIs were defined in each participant, six in the left (L) hemisphere and six in the right (R) hemisphere. These included six fROIs on the lateral surface of the frontal cortex in the inferior frontal gyrus (L/R IFG) and its orbital part (L/R IFGorb) as well as in the middle frontal gyrus (L/R MFG); and six fROIs on the lateral surface of the temporal and parietal cortex, in the anterior temporal cortex (L/R AntTemp), posterior temporal cortex (L/R PostTemp), and angular gyrus (L/R AngG).

For the MD network, eighteen fROIs were defined in each participant, nine in each hemisphere. These included the opercular part of the inferior frontal gyrus (L/R IFGop), the middle frontal gyrus (L/R MFG) and its orbital part (L/R MFGorb), the precentral gyrus (L/R PrecG), the insular cortex (L/R Insula), the supplementary motor area (L/R SMA), the inferior parietal cortex (L/R InfPar), the superior parietal cortex (L/R SupPar), and the anterior cingulate cortex (L/R AntCing).

For the DMN, ten fROIs were defined in each participant, five in each hemisphere. These included the posterior cingulate cortex (L/R PostCing), four medial frontal regions (L/R FrontMedOrb and L/R FrontMedSup), the precuneus (L/R Precuneus), and the temporoparietal junction (L/R TPJ).

### Examining the functional response profiles of fROIs

To estimate the responses of the fROIs to the conditions used to define them, we used an across-runs cross-validation procedure. In particular, for each relevant contrast (sentences > nonwords for the language regions, hard > easy for the MD regions, and easy > hard for the DMN regions), fROIs were defined for each participant based on data from only the first run, and their responses were then estimated using data from the second run. This procedure was then repeated using the second run to define the fROIs and the first run to estimate the responses. Finally, the responses were averaged across these two iterations to derive a single response magnitude for each condition in a given fROI/participant. This cross-validation procedure allows one to use all of the data for defining the fROIs as well as for estimating their responses (see Nieto-Castañón and Fedorenko, 2012, for discussion), while ensuring the independence of the data used for fROI definition and for response estimation (e.g., Kriegeskorte et al., 2009).

In order to compare the functional profiles across the three networks, we also estimated the responses of the fROIs to conditions that were not used to define them (i.e., hard and easy WM for the language network, sentences and nonwords for the MD network and DMN). Here, we used all of the data from the localizer task (i.e., both runs) to define the fROIs, and all of the data from the other task to estimate their responses.

Second-level analyses (repeated measures t-tests) were performed on these extracted response magnitude values, using false discovery rate (FDR) correction (Benjamini and Yekutieli, 2001) for the number of fROIs in each network. Pairwise comparisons of localizer effects across networks were tested using linear, mixed-effects regression models implemented with the lmer toolbox in R (Bates et al., 2015). These models included a fixed effect for network, a random slope of network by participant, and a random intercept by fROI. The fixed effect estimates were contrasted to each other using the “multcomp” package. Because these analyses were carried out to replicate previous findings, hypotheses were one-tailed.

### Examining inter-individual co-variation in effect sizes of the three networks

#### Descriptive statistics

For each functional contrast (sentences > nonwords, hard > easy WM, easy > hard WM) we computed, across participants, the Pearson correlations in effect size for every pair of fROIs (40 fROIs in total: 12 language fROIs, 18 MD fROIs, and 10 DMN fROIs). Then, the correlation values within each network and hemisphere (e.g., all pairwise correlations for regions of the left hemispheric language network) were Fisher-transformed and averaged together (this transformation reduces the bias in averaging correlations; see Silver and Dunlap, 1987). Correlations of fROIs with themselves, which were always equal to 1, were excluded from this step. Thus, we obtained: 3 networks × 2 hemispheres = 6 average correlations. Following a similar procedure, we computed the average inter-hemispheric correlation for each of the three networks (e.g., the average of all pairwise correlations between one left-hemisphere language region and one right-hemisphere language region). Similarly, we also computed the average correlation across pairs of networks, within each hemisphere (e.g., the average of all pairwise correlations between one left-hemisphere language region and one left-hemisphere MD region). Here, we obtained: 3 network pairs × 2 hemispheres = 6 average correlations. In total, the number of average correlations (within-network, within-hemisphere; within-network, across-hemispheres; and across-networks, within-hemispheres) was therefore 6+3+6 = 15.

For each of these 15 correlations we computed a 95% confidence interval via a bootstrapping procedure: first, we randomly sampled 5×10^4^ sets of n=60 participants, with replacement, from the observed data. Then, for each set, we re-computed the 15 correlations as outlined above, yielding – across all sets – 15 distributions of bootstrapped correlations. Finally, for each distribution, we found the interval that contained the middle 95% of values.

#### Significance tests

To test whether fROIs within a certain network *A* were more strongly correlated among themselves than with fROIs of another network B, we used a permutation approach. Specifically, we randomly shuffled effect sizes across participants for each fROI in A. Under this shuffling, the expected correlations within network *A* as well as between networks *A* and *B* are practically 0. Consequently, the difference between the mean pairwise correlation within *A* and the mean pairwise correlation between *A* and *B* is also expected to be 0. The effect of this shuffling procedure therefore corresponds to the null hypothesis that within-network correlations are no different from between-network correlations. Thus, to generate an empirical null distribution, shuffling is repeated many times (here, 5×10^−4^) and, for each repetition, the mean correlation between *A* and *B* is subtracted from the mean correlation within *A*. The resulting distribution is approximately normal, because of both the Fisher-transformation on pairwise correlations and the averaging of these correlations within / across networks. To test the probability that the observed data would be sampled under the null hypothesis, we fit a Gaussian to the null distribution and used its mean and standard deviation to *z*-score the observed data.

We used the same permutation approach to test the laterality of correlations in effect size between fROIs. Namely, we address three questions: (i) whether, for each network, correlations in effect size within each hemisphere are stronger than those across hemispheres; (ii) whether the size of this laterality effect differs across networks; and (iii) whether across-network correlations are also lateralized, such that fROIs in one network are differentially correlated with LH vs. RH fROIs in another network. The results of all of our tests are FDR-corrected for multiple comparisons (Benjamini and Yekutieli, 2001).

#### Hierarchical clustering

The permutation approach described above was used to test hypotheses-driven predictions regarding a tri-partite dissociation among the language, MD and default-mode networks. To use a more data-driven approach for examining our results, we searched for a partition of the 40 fROIs based solely on their pairwise correlations in effect size, ignoring their a-priori network labels. To this end, we first used a hierarchical clustering algorithm (Hartigan, 1975) to gradually connect all fROIs into a binary tree structure: this algorithm starts by joining the most correlated pair of fROIs into a node, and proceeds to join other pairs of fROIs and/or higher tree nodes in decreasing order of correlation (correlations between tree nodes are defined as the average pairwise correlation between their respective, constituent fROIs). Following the method of Blank *et al*. (2014), we then used a modularity-optimization approach to find the level at which the tree can be “ideally” partitioned into separate branches, each representing a group of fROIs; here, the “ideal” partition is one that maximizes within-branch correlations and minimizes across-branch correlations (a measure of “modularity”; see Gómez et al., 2009; Newman and Girvan, 2004).

## Results

### Behavioral data

Behavioral performance on the spatial working memory task was as expected: participants were more accurate and faster on the easy trials (accuracy=92.65±1.47%; reaction time (RT)=1.19±0.22s) than the hard trials (accuracy=79.81±2.39%, *t*_(59)_=-11.50, p<<10^−5^, Cohen’s *d*=1.48; RT=1.47±0.27s, *t*(59)=14.19, p<<10^−5^, *d*=1.83).

### Functional response profiles of the fROIs in the three networks

Replicating previous work, we find robust responses for all localizer contrasts using across-runs cross-validation (Figure 2). In the language network, the sentences > nonwords effect was highly reliable in each of the LH fROIs (*t*s>10.7, *p*s<0.0001, *d*s>1.38) and in each of the RH fROIs (*t*s>4.1, *p*s<0.0001, *d*s>0.53). In the MD network, the hard > easy effect was highly reliable in each fROI (*t*s>10.1, *p*s<0.0001, *d*s>1.30). And in the DMN network, the easy > hard effect was highly reliable in each fROI (ts>7.1, *p*s<0.0001, *d*s>0.92). Next, we examined the relationship between the language network and each of the other two networks:

#### Language vs. MD

Replicating prior work (Fedorenko, et al., 2011), we find no response to the spatial WM task in the language fROIs. None of the fROIs, except for the LMFG fROI, respond above baseline to either the hard spatial WM condition or the easy spatial WM condition (for both conditions, all *t*s<1, n.s., *d*<0.13). The LMFG fROI shows above baseline responses to both conditions, in line with what was reported in Fedorenko et al. (2011), and shows a slightly stronger response to the hard than the easy condition, which is not significant after an FDR correction for multiple comparisons (*t*_(59)_=1.84, n.s., *d*=0.24). In both hemispheres, the MD fROIs show an overall higher value for the hard > easy contrast than the language fROIs do (across-network contrast in LH: 0.74, *z*=7.8, *p*<10^−13^; RH: 0.66, SE=0.12, *z*=5.47, *p*<10^−7^).

Replicating Fedorenko et al. (2013), we find that the MD fROIs respond to the language localizer conditions in a manner opposite to the language fROIs. In particular, they respond more to the meaningless and unstructured nonword lists than to the sentence condition (*t*s>2.1, *p*s<0.02, *d*s>0.27). (Interestingly, this pattern holds even though the task in the current version of the language localizer is passive reading (cf. a memory probe task in Fedorenko et al., 2013); see also Fedorenko (2014).)

#### Language vs. DMN

In line with much prior work (e.g., Gusnard and Raichle, 2001; Raichle et al., 2001), we find that the DMN fROIs de-activate to the spatial WM task, with both the hard and the easy condition eliciting a response reliably below the fixation baseline (hard: *t*s>7.1, *p*s<10^−9^, *d*s=0.92; easy: *t*s>4.2, *p*s<10^−4^, *d*s>0.54). Critically, in sharp contrast with the language fROIs, none of the fROIs, except for the LTPJ fROI, respond above baseline to sentence comprehension (sentence condition: *t*s<1.1, n.s., *d*s<0.14). The LTPJ fROI responds above baseline to the sentence condition (*t*_(59)_=4.56, *p*<10^−4^, *d*=0.59) and reliably more to sentences than nonwords (*t*_(59)_=5.14, *p*<10^−5^, *d*=0.66). Even so, this response to the sentence condition is ~3.5 times lower than the response in the nearby LPostTemp language fROI. Directly comparing the overall response to the sentence condition across the language network and DMN showed that the responses in the former are significantly higher (across-network contrast in LH: 1.31, *z*=7.99, <10^−14^; RH: 0.67, *z*=5.08, *p*<10^−6^). Therefore, although several language fROIs show a hint of the DMN signature (deactivation to the demanding spatial WM task, with stronger deactivation to the harder condition, consistent with: Fedorenko et al., 2011), the functional response in the DMN fROIs is clearly distinct from that in the language fROIs: language, but not DMN, fROIs respond robustly during language comprehension.

### Within-vs. between-network correlations in effect size

The correlations of effect sizes for all pairs of fROIs are shown in Figure 3. Even before performing any statistical analysis, the dissociation among the language, MD, and default mode networks is visually apparent; the correlations are much higher for pairs of regions within each network than for pairs of regions across networks. A quantitative summary of the effect-size correlations is shown in Figure 4.

In line with previous work, we find a dissociation between the language and MD networks. In the left hemisphere, the average correlation between the sentences > nonwords effect size in one language region and the sentences > nonwords effect size in another language region is 0.54 (95% confidence interval (CI) = [0.39, 0.67]), similar to what was reported in Mahowald & Fedorenko (2016). In other words, individuals who show a bigger (smaller) sentences > nonwords effect in a given language region also tend to show bigger (smaller) effects in other language regions. Similarly, the average correlation between the hard > easy effect size in one MD region and the hard > easy effect size in another MD region is 0.66 (CI_95_%=[0.55, 0.76]). However, the average correlation between the sentence > nonwords effect size in a language region and the hard > easy effect size in a MD region is 0.18 (CI_95%_=[0.03, 0.31]), significantly lower than both the within-language (*p*<10^−13^) and within-MD (*p*≈0) correlations. In other words, individual differences in the size of the sentences > nonwords effect in the language network and the size of the hard > easy effect in the MD network are less predictive of one another, relative to individual differences among fROIs within each network.

**Figure 2.**
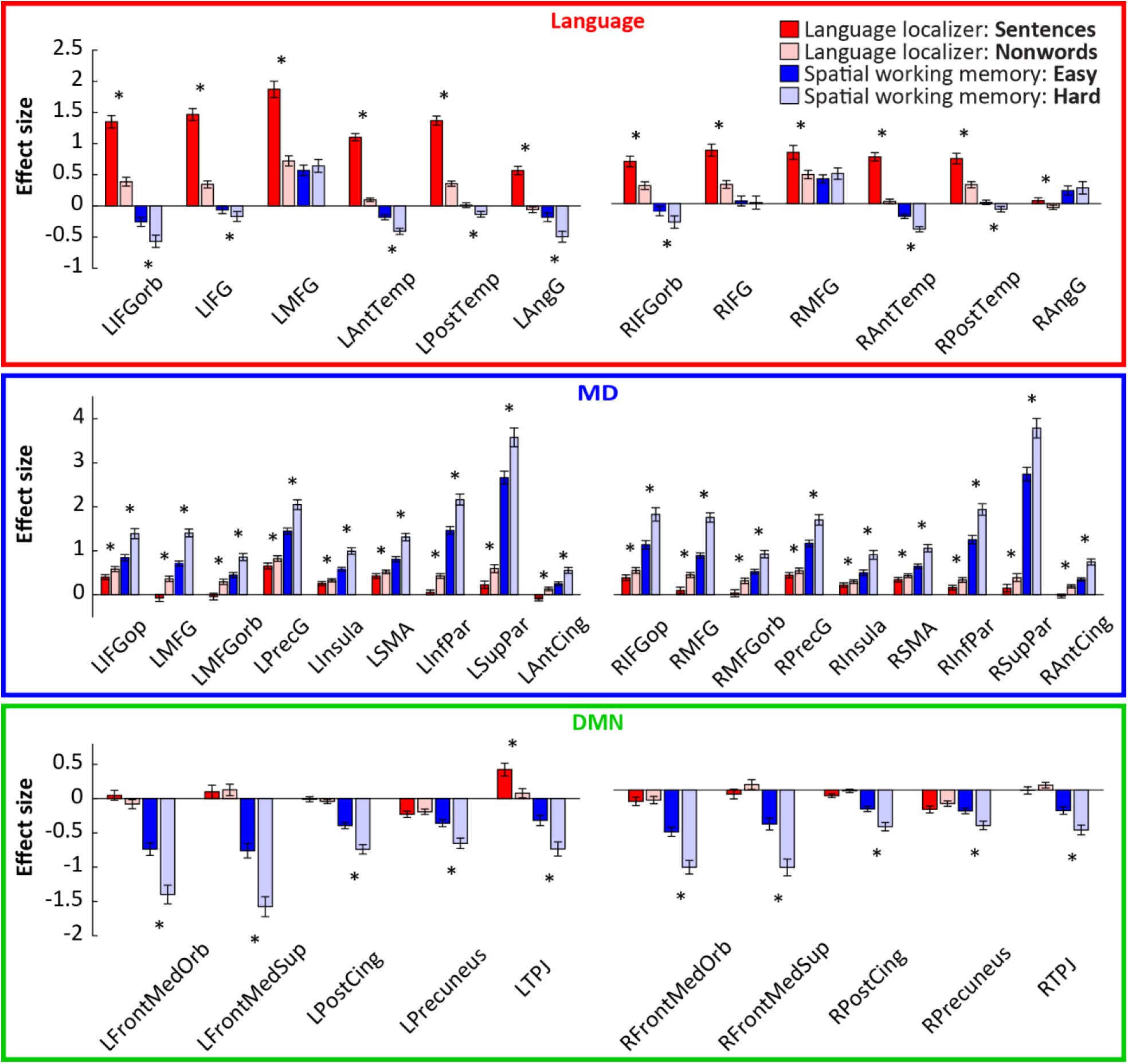
Responses to the conditions of the localizer tasks in each fROI. The language fROIs (top) are defined by the sentences > nonwords contrast; the MD fROIs (middle) are defined by the hard > easy spatial working memory contrast; and the DMN fROIs (bottom) are defined by the easy > hard spatial working memory contrast. The responses to the conditions used for defining the fROIs are estimated using across-runs cross-validation, to ensure independence. Left: left-hemispheric fROIs (L prefix). Right: right-hemispheric fROIs (R prefix). Significant effects (after an FDR-correction for multiple comparisons within each network) are marked with a black star. IFG: inferior Frontal Gyrus; IFGorb: IFG *pars orbitalis*; MFG: middle frontal gyrus; AntTemp: anterior temporal cortex; PostTemp: posterior temporal cortex; AngG: angular gyrus; IFGop: IFG *pars opercularis*; MFGorb: MFG, orbital part; PrecG: precentral gyrus; SMA: supplementary motor area; InfPar: inferior parietal cortex; SupPar: superior parietal cortex; AntCing: anterior cingulate cortex; FrontMedOrb: medial frontal cortex, orbital part; FrontMedSup: medial frontal cortex, superior part; PostCing: posterior cingulate cortex; TPJ: temporo-parietal junction.

**Figure 3.**
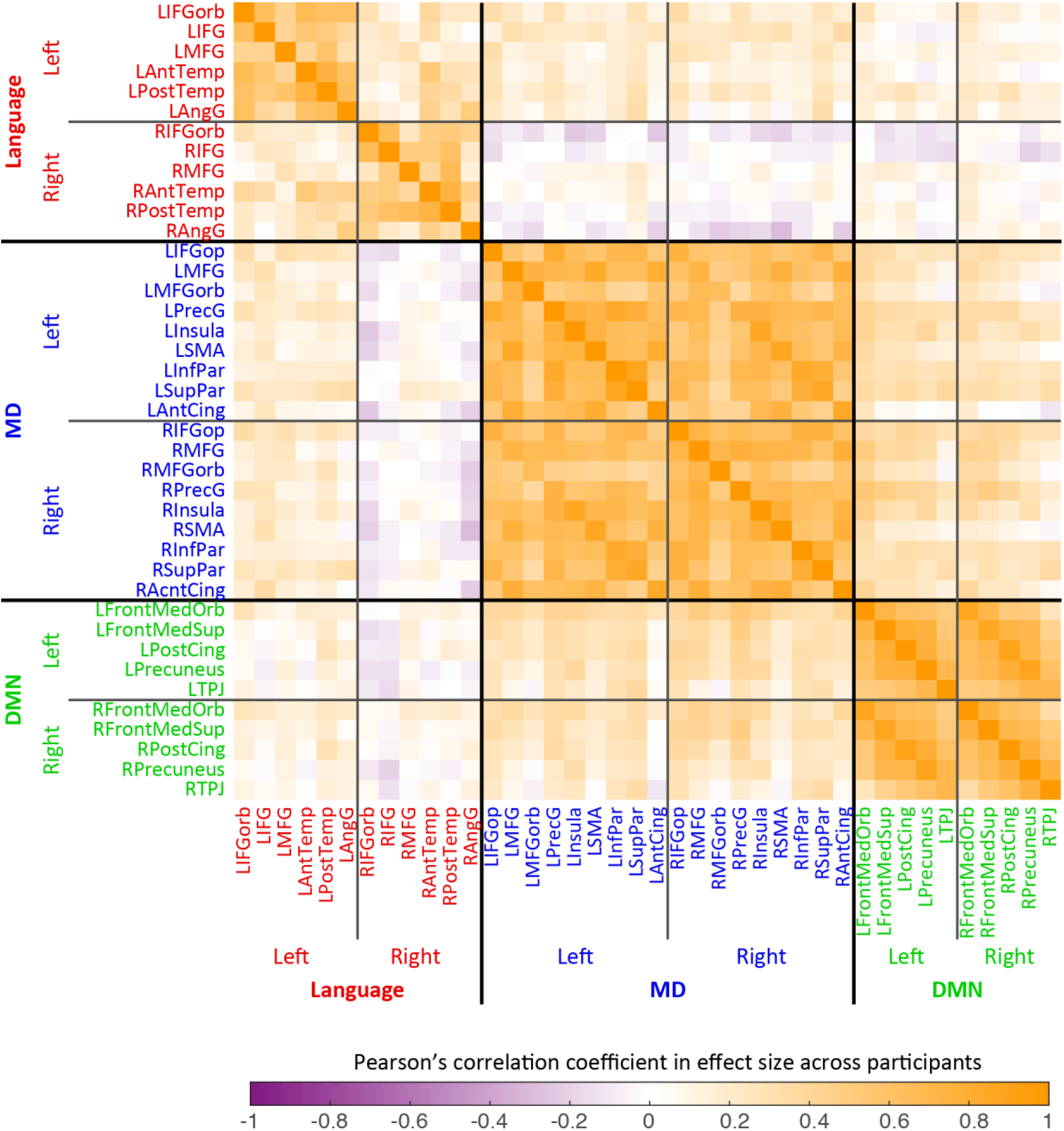
Pearson correlations, across 60 participants, for effect sizes of localizer contrasts across pairs of fROIs. For each fROI, the effect size is for the contrast used to define that fROI (but estimated in independent data): for the 12 language regions (labeled in red font), the sentences > nonwords contrast from the language localizer task was used; for the MD regions (blue font), the hard > easy spatial WM contrast was used; and or the DMN regions (green font), the easy > hard spatial WM contrast was used.

Critically, in line with the distinct functional profiles reported above, we also observe a clear dissociation between the language network and the DMN. The average correlation between the sentences > nonwords effect size in a language region and the easy > hard effect size in a DMN region is 0.09 (CI_95_%=[-0.08, 0.26]). This is significantly lower than the within-language correlations (*p*≈0) and the within-DMN correlations (0.66, CI_95%_=[0.58, 0.74]; *p*≈0). The DMN is also dissociated from the MD network, with between-network correlations in effect size (0.26, CI_95%_=[0.08, 0.43]) significantly lower than those within each network (both *p*≈0).

A similar dissociation among the three networks obtains in the right hemisphere, as shown in Figures 3 and 4: the mean pairwise correlations within the language network (0.47, CI_95%_=[0.36, 0.58]) and within the MD network (0.65, CI_95%_=[0.54, 0.75]) are stronger than the mean language-MD pairwise correlation (-0.05, CI_95%_=[-0.18, 0.11]; both p≈0); the correlations within the language network and within the DMN (0.64, CI_95%_=[0.54, 0.74]) are stronger than language-DMN correlations (0.04, CI_95%_=[-0.10, 0.17]; both *p*≈0); and the correlations within the MD network and within the DMN are stronger than MD-DMN correlations (0.25, CI_95%_=[0.09, 0.40]; *p*≈0 and *p*<10^−13^, respectively).

Furthermore, the language network shows a robust lateralization effect in this novel measure, such that LH language fROIs are more correlated among themselves than they are with RH language fROIs (mean inter-hemispheric correlation: 0.22, CI_95%_=[0.03, 0.40]; *p*<10^−12^), and the same is true for correlations among RH language fROIs (*p*<10^−6^). These findings are in line with prior functional correlation studies (e.g., Blank et al., 2014; Gotts et al., 2013) and dynamic network modeling studies (e.g., Chai et al., 2016). In contrast, this laterality effect is not observed in either the MD network (mean inter-hemispheric pairwise correlation: 0.62, CI_95%_=[0.50, 0.73]; compared to LH correlations, *p*=0.40; compared to RH correlations, *p*=0.97) or the DMN (mean inter-hemispheric correlation: 0.68, CI_95%_=[0.58, 0.76]; compared to LH correlations, *p*=1; compared to RH correlations, *p*=0.72). A direct comparison of laterality effects across networks further confirms that they are stronger in the language network than in either the MD or the DMN, for both the LH (both *p*<10^−5^) and the RH (MD: *p*=0.001; DMN: *p*<10^−5^). A related, potentially interesting observation is that MD fROIs are more correlated with the LH language fROIs than they are with the RH homologues (LH MD fROIs: *p*=0.01; RH: *p*=0.003). In all other cases, across-network correlations in effect-size do not show such laterality.

**Figure 4.**
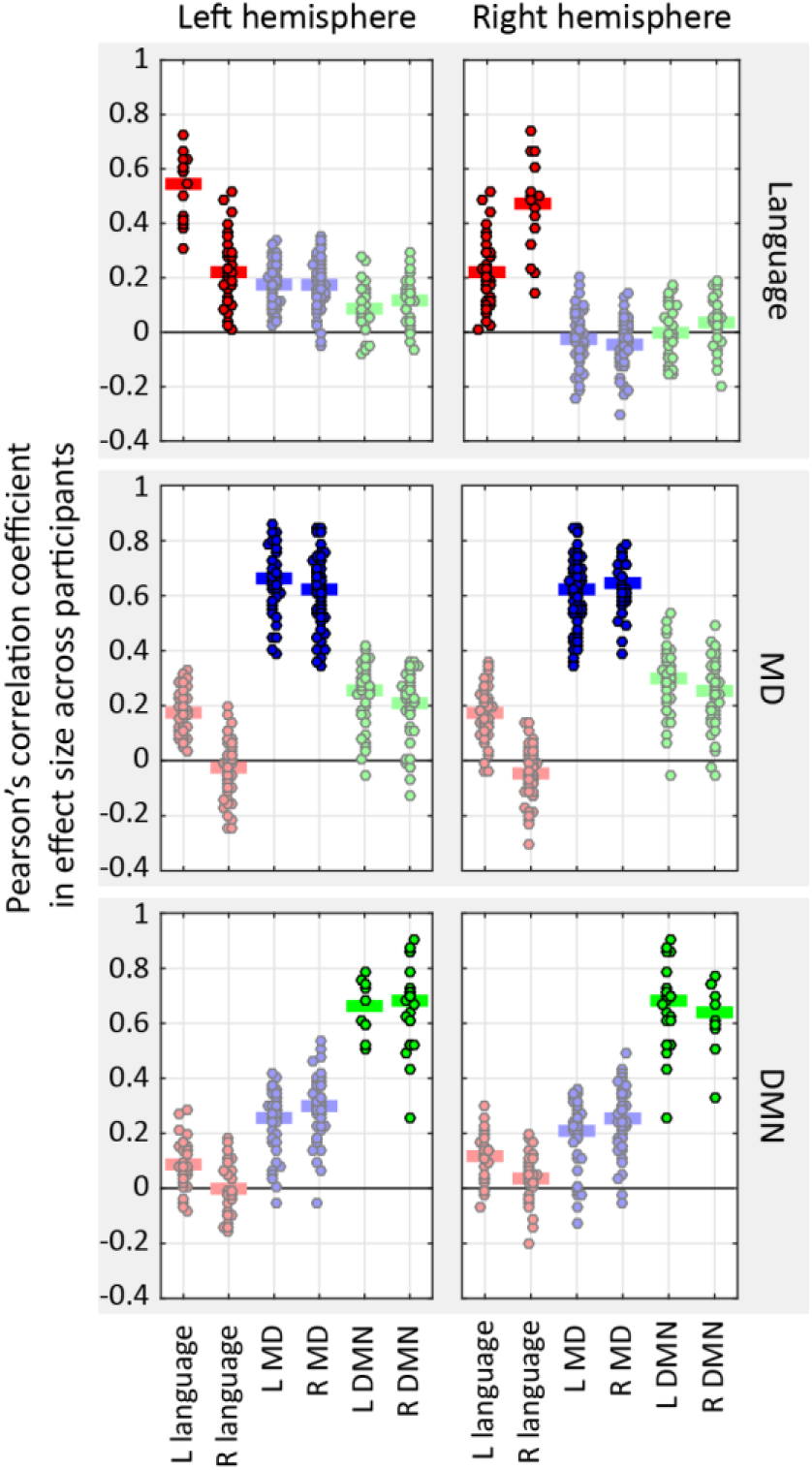
Correlations in effect sizes across participants, computed either within networks (strong colors) or between networks (faint colors). Each point is a pairwise correlation between the effect sizes of two fROIs, one from the network/hemisphere denoted on the *x*-axis, and one from the network/hemisphere denoted by the subplot titles (top: language; middle: MD; bottom: DMN; left: left hemisphere; right: right hemisphere). Horizontal lines show averages across these pairwise correlations.

These findings, revealed via hypothesis-driven tests in which sets of fROIs are compared to each other based on a pre-determined division into functional networks, are also supported by a hypothesis-neutral, data-driven analysis (Figure 5). Namely, hierarchically clustering all fROIs into a tree-structure based on their pairwise correlations in effect size – without a-priori information on their network assignments – recovered the dissociation among language, MD and DMN fROIs as well as the associated laterality patterns. Specifically, the tree obtained from this clustering contained four branches precisely corresponding to the LH language, RH language, bilateral MD, and bilateral DMN networks. This partition had the highest modularity value compared to all other partitions, both finer and grosser, that were licensed by the tree – indicating that it was the overarching organizing principle in our data. (In contrast to the branching of language fROIs by hemisphere, the MD and DMN branches were overall organized by inter-hemispheric homology such that many fROIs clustered with their respective contra-lateral homologues before forming larger clusters with one another.)

**Figure 5.**
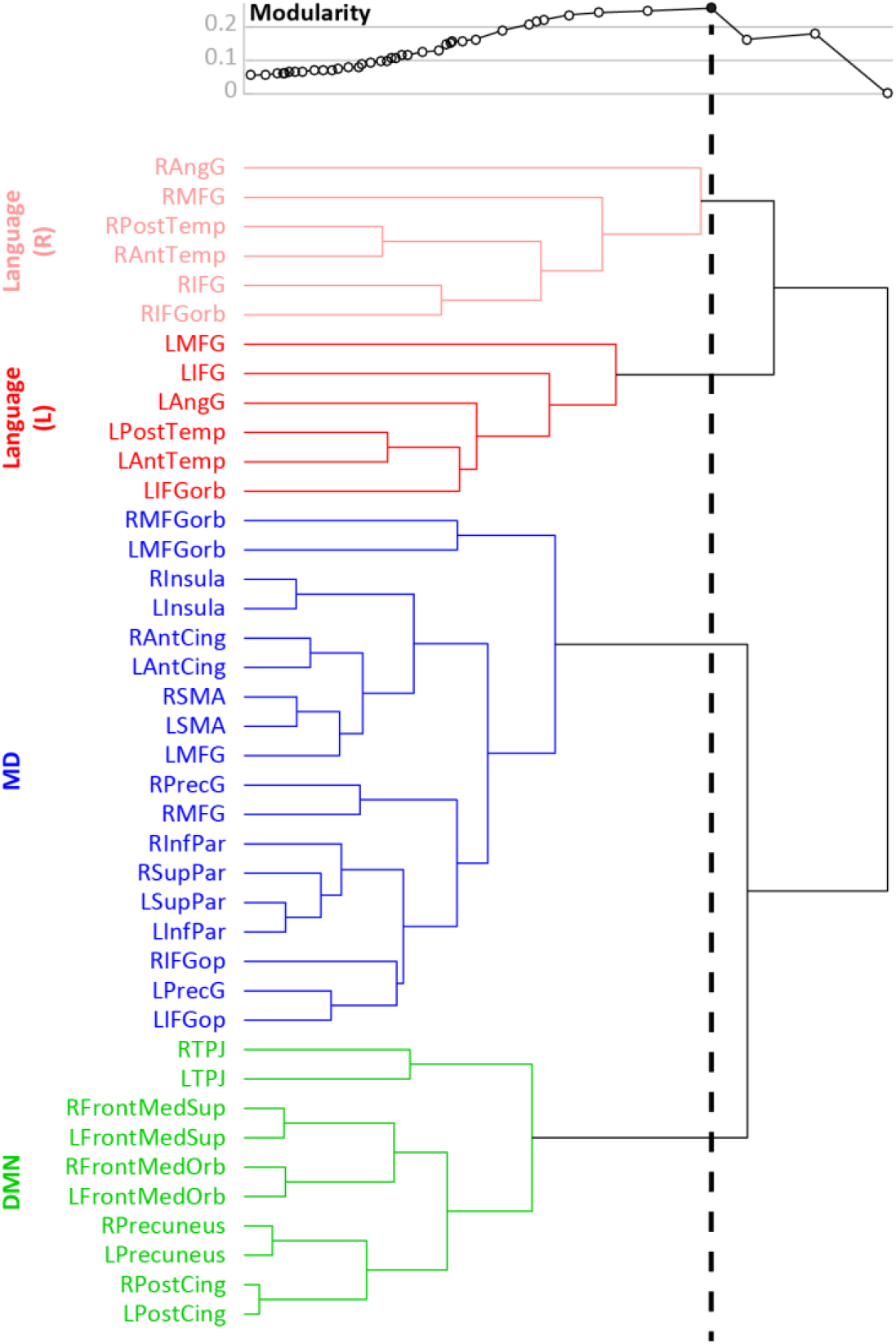
Hierarchical clustering results. In the binary tree shown, branch length (i.e., horizontal lines) corresponds to the similarity between fROIs (or sets of fROIs). Above the tree, modularity is plotted for all fROI partitions licensed by the tree. Each point on the modularity plot corresponds to a partition generated by drawing an imaginary vertical line from that point through the tree and clustering together only those fROIs that are merged to the left of this line (fROIs that are merged to the right of the line remain in separate clusters). A sample vertical line is drawn for the maximal modularity.

The results reported above are all based on correlations between effect sizes from the language localizer task (in language fROIs) and effect sizes from the spatial working-memory task (in MD and DMN fROIs). Differences between these two tasks might therefore trivially account for the functional dissociation among the three networks. In order to reject this account, we re-computed our critical measure – i.e., inter-region correlations in effect size across participants – based on data from a single localizer contrast across all fROIs. Specifically, we measured the sentences > nonwords effect size in each language, MD and DMN fROI and, then, compared the average inter-regional correlation within the language network to the average language-MD correlation and the average language-DMN correlation. Similarly, we measured the hard > easy (or easy > hard) effect size in each fROI and, then, compared the average interregional correlation within the MD (DMN) network to the corresponding inter-network correlations. The results of this analysis are presented in Table 1. Consistent with our main analysis, they indicate a clear tri-partite dissociation across the three networks.

**Table 1.** Mean inter-regional correlations in contrast-specific effect sizes across participants^a^

| Localizer contrast | *r* |  |
| --- | --- | --- |
|  | LH | RH |
| *Reading: Sentences > Nonwords* |  |  |
| Within language | 0.54 | 0.47 |
| Language vs. MD | 0.10 | 0.16 |
| Language vs. DMN | 0.30 | 0.09 |
| *Spatial WM: Hard > Easy* |  |  |
| Within MD | 0.66 | 0.65 |
| MD vs. language | 0.10 | 0.13 |
| MD vs. DMN | -0.26 | -0.25 |
| *Spatial WM: Easy > Hard* |  |  |
| Within DMN | 0.66 | 0.64 |
| DMN vs. language | 0.38 | 0.25 |
| DMN vs. MD | -0.26 | -0.25 |

<sup>a</sup> All within-network correlations are stronger than their respective across-network correlations at $p < 10^{-7}$

## Discussion

The current study examined the relationship among three large-scale functional networks that support high-level cognitive processes: the language network, the multiple demand (MD) network, and the default mode network (DMN). To do so, we (a) characterized the functional response profiles of each network, and (b) employed a novel analytic approach that tested, across participants, the correlations in response magnitude among the fROIs within each network vs. between networks. Using both analyses, we replicate the dissociation between the language and MD networks (e.g., Blank et al., 2014; Fedorenko et al., 2011; Fedorenko et al., 2013; Paunov et al., submitted), as well as the well-established dissociation between the MD network and the DMN. Critically, we further demonstrate that the language network is also robustly dissociable from the DMN: the former, but not the latter responds strongly during language processing and, whereas regional effect sizes strongly co-vary across individuals within each network, there is little or no such correlation between the two networks. In other words, if an individual shows a strong response to language processing (the functional signature of the language system) in one language region, they will also show a strong response in other language regions. Similarly, if an individual shows strong deactivation to a demanding task (the functional signature of the DMN) in one DMN region, they will also show strong deactivation in other DMN regions. However, the strength of the response to language processing in a language region bears little information on how much a DMN region will deactivate (or how much a region of the MD network will respond) to a demanding task.

These results have two implications. The first one is methodological: inter-individual differences in effect sizes do not simply reflect general variability in either brain structure (e.g., vascularization affecting the fMRI BOLD signal) or in behavioral/cognitive states (e.g., attention). If this were the case, we would expect effect sizes in different brain regions to strongly co-vary across individuals, regardless of their functional profiles. Instead, such interindividual differences appear to be sensitive to the functional architecture of the brain, respecting its division into distinct, large-scale neural networks. Thus, inter-region correlation in effect size across individuals is a powerful new measure for discovering functional dissociations among neural systems and, possibly, even at a finer grain within each system.

The second implication is theoretical. The prior literature has left the relationship between the language network and the DMN ambiguous. In particular, methods that cluster voxels across the brain based on their respective activity time-courses sometimes recover a network that looks like a combination of the language network and the DMN (e.g., Yeo et al., 2011); this result appears to depend, in part, on the pre-specified number of clusters that such analyses are constrained to produce. Further, although the language regions show no response to non-linguistic demanding tasks (and are thus clearly dissociable from the domain-general MD network), they sometimes show deactivation to such tasks, much like the DMN (e.g., Fedorenko et al., 2011; see also Figure 2, top panel). Finally, both the language and the DMN regions have been linked to semantic / conceptual processing (e.g., Binder et al., 2009; Jackson et al., 2016; Wirth et al., 2011). However, we find a clear and robust functional dissociation between the language network and the DMN. This finding suggests that, in spite of some functional similarities between these two networks, and in spite of the fact that some of their regions lie in close proximity to one another, these two networks likely support distinct computations. Thus even if both networks get engaged in the service of the same goal (supporting some aspects of conceptual processing; e.g., Binder et al., 2009), they plausibly differ in their respective contributions and should be treated as cognitively separable when hypotheses about their functions are evaluated.

## Acknowledgments

This work was supported by award HD 057522 to E.F. from NICHD, and by a grant from the Simons Foundation to the Simons Center for the Social Brain at MIT. K.M. was supported by a National Defense Science and Engineering Graduate (NDSEG) Fellowship. We would like to acknowledge the Athinoula A. Martinos Imaging Center at McGovern Institute for Brain Research at MIT, and the support team (Steve Shannon, Atsushi Takahashi, and Sheeba Arnold).

